# Degraded RNA from human anterior cruciate ligaments yields valid gene expression profiles

**DOI:** 10.1101/2022.07.14.500070

**Authors:** Matthew D. Strawn, Geoffrey D. Fisher, Cody J. Perry, Javad Hashemi, Hossein Mansouri, Matthew P. Ferguson, Mimi Zumwalt, George W. Brindley, James R. Slauterbeck, Daniel M. Hardy

## Abstract

**Background:** Correlating gene expression patterns with biomechanical properties of connective tissues provides insight into molecular processes underlying development and maintenance of the tissues’ physical characteristics. Cadaveric tissues such as human knees are widely considered suitable for biomechanical studies, but their usefulness for gene expression experiments is potentially limited by unavoidable, nuclease-mediated degradation of RNA.

**Methods:** We quantified mRNAs encoding 84 extracellular matrix components and five housekeeping proteins in intact and partially degraded RNA from human anterior cruciate ligaments (ACLs), and compared relative amounts of the individual mRNAs by regression analysis.

**Results:** Human ACL RNA degraded *in vitro* by limited ribonuclease digestion (N=6) resembled degraded RNA isolated from cadaveric tissue. PCR threshold cycle (Ct) values from degraded RNAs ranged variably higher than values obtained from their corresponding non-degraded RNAs, reflecting both the expected loss of target templates in the degraded preparations as well as differences in the extent of degradation. Relative Ct values obtained for mRNA targets in degraded preparations correlated strongly with the corresponding target levels in non-degraded RNA, both for each ACL as well as for pooled results from all six ACLs. Nuclease-mediated degradation produced similar, strongly correlated losses of housekeeping and non-housekeeping gene mRNAs. Expression profiling of RNA degraded *in situ* yielded comparable results, confirming that *in vitro* digestion suitably mimicked degradation by endogenous ribonucleases in frozen and thawed ACL.

**Conclusion:** PCR-based expression analyses can yield valid mRNA profiles from partially degraded RNA preparations such as those obtained from cadaveric knees and other skeletal motion segments used in biomechanical studies. Legitimate comparisons between variably degraded tissues can be made by normalizing quantitative data to an appropriate housekeeping transcript.

## INTRODUCTION

Profiles of expressed mRNAs reflect the physiological states of cells and tissues ^1,2^. Consequently, comparative expression profiling can identify genes and metabolic pathways associated with onset, progression, and treatability of disease ^3,4^. Approaches to mRNA profiling range from global analyses such as RNA-Seq or microarray screening of all expressed genes, to more focused, PCR-based profiling of gene subsets. Regardless of the approach, expression profiling methods universally call for use of only high quality, intact RNA to ensure that results accurately reflect the pattern of mRNAs present in living cells or tissues. Appropriate methods for tissue acquisition and handling and for RNA extraction and purification can yield intact, high quality RNA even from difficult tissues such as pancreas and ligament^5,6^. However, RNA degradation prior to isolation is sometimes unavoidable in precious specimens such as human cadaveric tissues that cannot always be handled and stored under ideal conditions, thus potentially limiting the usefulness of RNA from these tissues for expression analyses.

RNA profiling studies have characterized molecular responses of connective tissues to injury in non-human animals ^6,7^, and identified sex differences in gene expression that may underlie susceptibility of female athletes to anterior cruciate ligament (ACL) injury ^8,9^. However, RNA integrity often cannot be preserved completely in biomechanical studies of human cadaveric tissues, which confounds studies seeking, for example, to correlate gene expression with the human ACL’s ultrastructure and biomechanical properties (refs. 10, 11, and unpublished observations), or to identify expression patterns associated with degeneration and biomechanical properties of human intervertebral discs (Fisher et al., unpublished). Some studies have shown that meaningful expression data can be obtained using degraded RNA from sources such as rectal tumors ^12^ or cultured cells ^13^, but none have assessed validity of profiles generated using degraded RNA from human connective tissues commonly used for biomechanical studies.

Here, we tested the hypothesis that gene expression profiles generated using partially degraded RNA from human ACLs accurately reflect those generated using intact RNA.

### METHODS

### Tissue acquisition

We recovered human ACL tissues, by IRB-approved protocol (TTUHSC IRB# L06-154), during total knee arthroplasty or ACL reconstruction surgeries (8 male, 5 female; ages 29-73 years, mean 62.3 years), and immediately froze [N_2_ *(l)*] the tissues for storage at -80 °C. For human intervertebral disc tissue, we excised the L4-5 disc from the frozen spine of a willed body (non-embalmed female, age 40; spine stored at -20°C). As is commonplace for donated bodies, no information was available regarding the time between donor death and freezing of the spine, or the specific conditions of its storage prior to receipt in the lab.

### RNA isolation

We isolated RNA from frozen [N_2_ *(l)*], thoroughly pulverized tissues (whole ACL or the outer anterior segment of the intervertebral disc’s annulus fibrosus) using either the RNeasy Mini Kit (100-350 mg of tissue; QIAGEN, Valencia, CA) or the guanidinium thiocyanate/acidic phenol extraction^8^ method (1 g tissue) and assessed RNA purity and concentration by spectrophotometry (NanoDrop™, Thermo Fisher Scientific, Wilmington, DE). RNA degradation. For *in vitro* degradation of ACL RNA, we digested paired aliquots (4 µg, isolated by RNeasy kit) either with RNase ONE™ (0.005 units, 37 °C, 1 h; Promega, Madison, WI) to produce ‘partially degraded’ RNA, or with no added enzyme to produce a mock-digested, ‘intact’ control RNA, and assessed extent of degradation by formaldehyde/agarose gel electrophoresis (1 µg RNA/lane) ^8, 14^.

For *in situ* degradation of ACL RNA, we incubated 1 g of thawed, pulverized ACL for 20 min at 23°C to permit autolytic degradation of RNA by endogenous RNAses, isolated RNA by the guanidinium thiocyanate/acidic phenol extraction method^8^, then assessed RNA quality with an Agilent Technologies 2100 Bioanalyzer (Santa Clara, CA). A matching 1 g portion of unthawed, pulverized ACL served as a non-degraded control.

### Quantitative PCR

We used first-strand complementary DNA (cDNA) synthesized from 500 ng of RNA with combined oligo-dT and random hexamer priming (RT2 First Strand Kit; SABiosciences, Frederick, MD) as template for real time PCR (Human Extracellular Matrix and Adhesion Molecules RT2 Profiler™ PCR Array kit; SABiosciences), with SYBR Green detection. We calculated threshold cycle (Ct) values at threshold=0.4 from amplification plots (ABI Prism 7000 Sequence Detection System; Applied Biosystems, Foster City, CA), and omitted from further analysis any genes the software identified as “undetermined” because they did not amplify sufficiently during the exponential phase. All five housekeeping genes in the array amplified successfully in all profiles, so downstream correlation analyses (see below) included Ct values from the full complement of these genes.

### Data analysis

To determine the effect of RNA degradation on expression profile, we correlated C_t_ values obtained from degraded RNAs and non-degraded RNAs using hierarchical regression models. We first established a linear relationship by analyzing all data from the six ACL RNA preparations, then produced individual regression equations for ACL RNAs by treating them as random samples from the population regression, and examined the assumption of randomness of the regression coefficients by fitting hierarchical linear regressions with random slopes and random intercepts. To analyze variation in degradation, we averaged the differences in Ct values between intact and degraded RNAs for each gene in each of the six ACLs, then plotted the averages for housekeeping and non-housekeeping genes against each other to test for correlation. Finally, we performed two-tailed t-tests to identify differences in extent of degradation between the housekeeping and non-housekeeping groups.

## RESULTS

Limited RNase digestion of ACL RNAs yielded degraded preparations with electrophoretic properties similar to partially degraded RNA isolated from human cadaveric intervertebral disc (Fig. 1). Notwithstanding identical digestion conditions, the six degraded ACL RNA preparations exhibited subtle electrophoretic differences, with RNAs from ACL #98 and #104 appearing slightly less degraded than the other four (Fig. 1). Using hierarchical models, regression analysis of all C_t_ value pairs (degraded vs. intact) obtained from the six ACL RNAs revealed a strong linear relationship (Pearson correlation coefficient r=+0.80) given by the equation

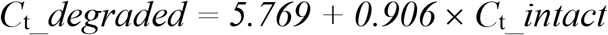

**Fig. 1.**
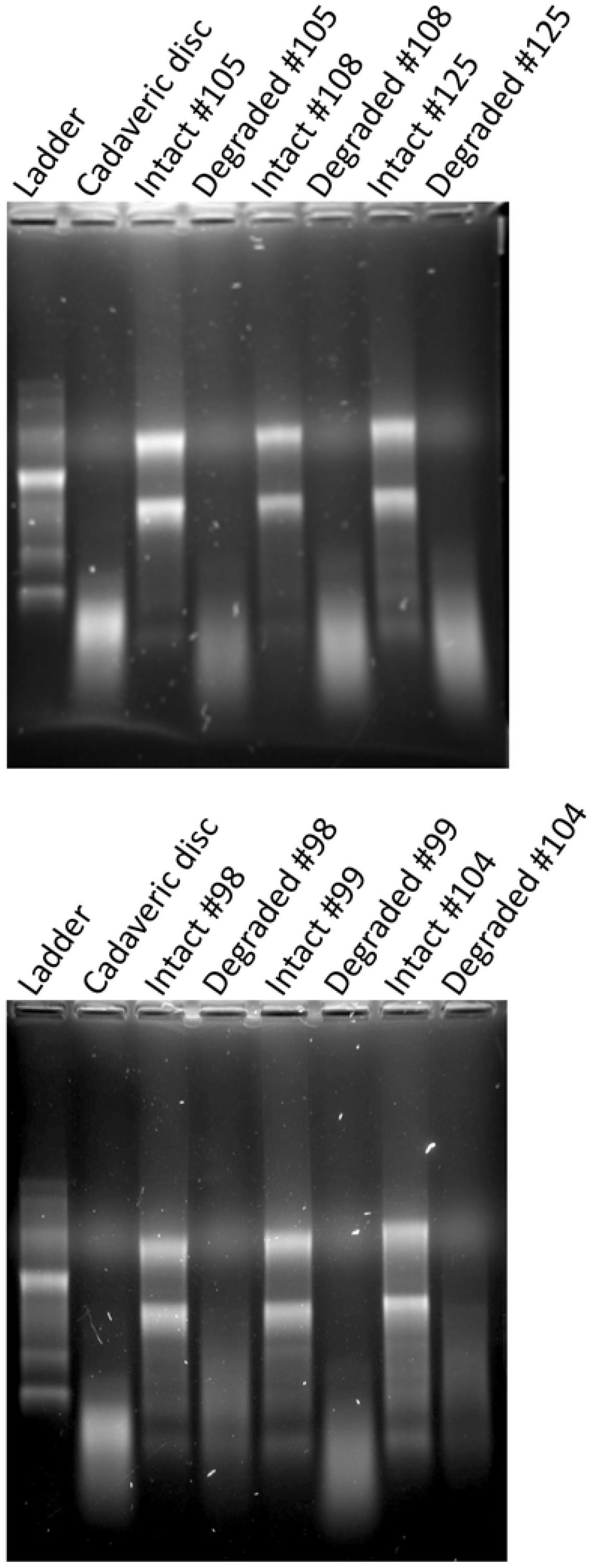
Formaldehyde-agarose electrophoresis of intact and partially degraded RNA preparations. We evaluated paired samples of intact and *in vitro*-degraded RNA from six ACLs (designated #105, 108, 125, 98, 99, and 104). Note the loss of ribosomal RNA bands and appearance of smaller-sized RNA fragments upon RNAse digestion of the ACL RNAs, as well as the similarity of the electrophoretic patterns to the degraded RNA isolated from cadaveric intervertebral discs.

The line intercept (95% confidence interval *3*.*254 - 8*.*284* cycles) reflected an increase in the number of PCR cycles required to reach the threshold signal as a consequence of degradation-induced decreases in the amounts of amplifiable mRNAs for the 89 targets. The line slope near but slightly below 1.0 (95% confidence interval: *0*.*868 – 0*.*943*) revealed that degradation produced a generally uniform increase in C_t_ irrespective of an individual RNA’s abundance, with a possible trend toward proportionally greater loss of higher abundance mRNAs.

Hierarchical analysis of individual regression lines for the six RNA preparations (see Fig. 2 for example individual regression plots) revealed strong linear relationships (range of Pearson coefficients: 0.86-0.98; Table 1), with non-random slopes of the six lines each fixed at 0.906 as for the population overall. In contrast, randomly distributed intercepts ranged from 2.93 – 7.48, reflecting possibly stochastic variation in the extent of RNA degradation between preparations. Mean Ct increase ranged from 0.75 cycles for the 86 targets detected in degraded ACL 104 RNA, to 5.2 cycles for the 80 targets detected in degraded ACL 105 RNA, corresponding to 40% and 97% average losses of amplifiable target, respectively [fraction remaining = *f* = 2exp(−1’C_t_); % loss = (1 – *f*)×100]. The randomly distributed y-intercept (b) values correlated inversely with the number of targets quantified (N) and with magnitude of the Pearson coefficient (r), indicating that as average C_t_ increased because of RNA degradation, more targets were lost and the scatter of the data increased. Nonetheless, even at the highest levels of degradation (ACLs 99, 105, 108, and 125), profiles from individual degraded RNAs correlated strongly with those from the corresponding intact RNAs (r > +0.86). Likewise, the Coefficients of Determination (R^2^) ranging from 0.75-0.96 indicated that 75-96% of the variation in target signal from degraded RNAs reflected the biologically relevant distribution of target signals from the corresponding intact RNA.

**Fig. 2.**
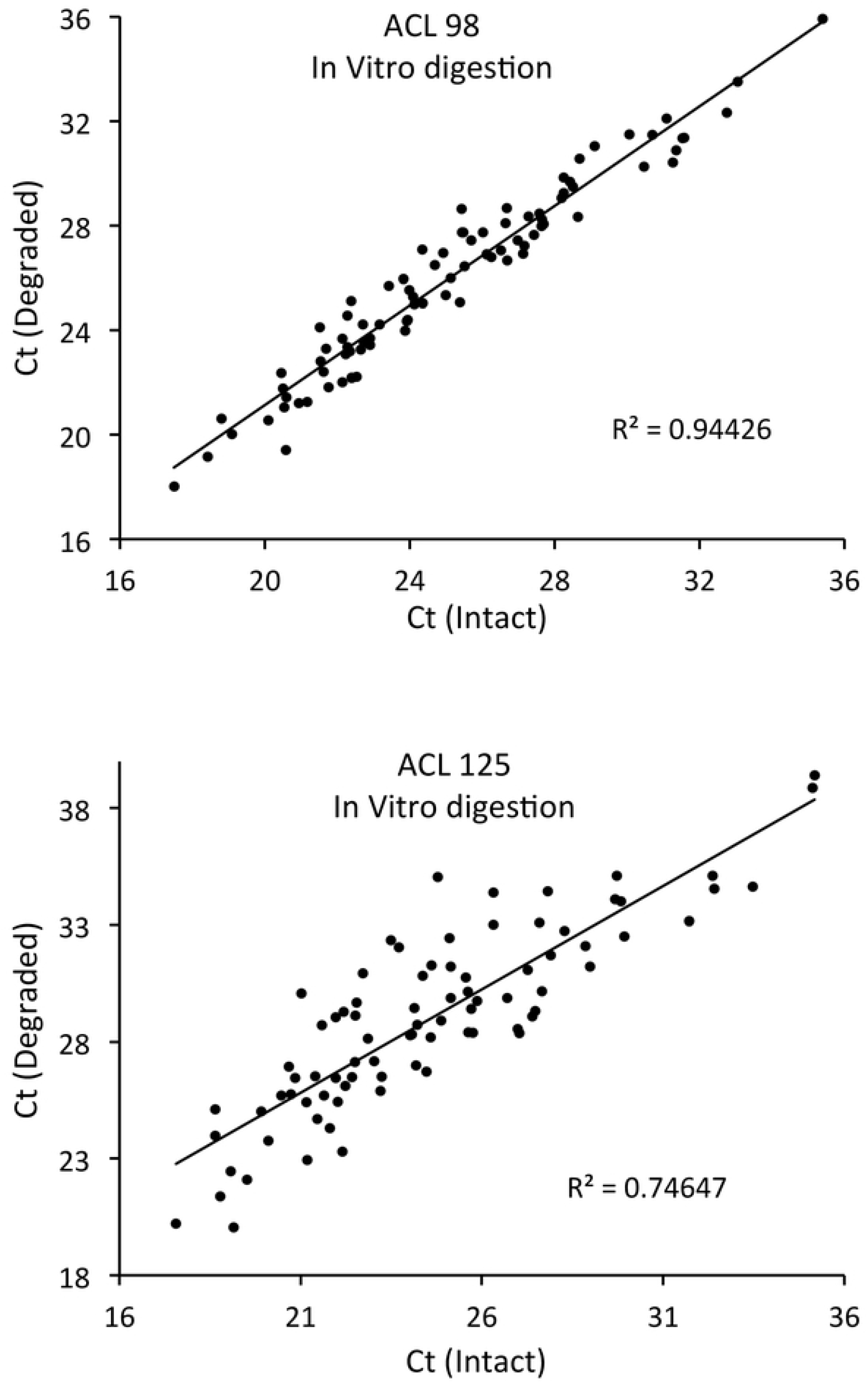
Example linear regression plots of Ct values from intact vs. *in vitro*-degraded RNA for two ACLs (#98 and #125) representing relatively less or more degradation, respectively (compare to Fig. 1). See Table 1 for a summary of results from corresponding analyses of degraded RNA preparations from all six ACLs.

**Table 1.** Correlation of mRNA expression profiles from partially degraded ACL RNAs with profiles from their corresponding intact RNAs. Differences in N (number of genes compared) reflect differences in the number of C_t_ values returned for each ACL, with N=89 representing no “undetermined” targets for an individual ACL owing to degradation of RNA target template to a level below that required for PCR amplification. Note that all six ACL RNAs yielded lines with slope 0.906, the same as for the combined data, but with randomly distributed y-intercept values (b). Note also the higher Coefficients of Determination, ranging up to a very strong positive correlation of 0.96, for all individual lines in comparison to the combined data.

| ACL # | Targets Compared (N) | Slope (m) | y-intercept (b) | Pearson Coefficient (r) | Coefficient of Determination ($R^2$) |
| --- | --- | --- | --- | --- | --- |
| Combined | 497 | 0.906 | 5.77 | +0.80 | 0.64 |
| 98 | 87 | 0.906 | 3.28 | +0.97 | 0.94 |
| 99 | 83 | 0.906 | 7.19 | +0.88 | 0.77 |
| 104 | 86 | 0.906 | 2.93 | +0.98 | 0.96 |
| 105 | 80 | 0.906 | 7.48 | +0.87 | 0.76 |
| 108 | 78 | 0.906 | 7.01 | +0.88 | 0.77 |
| 125 | 83 | 0.906 | 6.72 | +0.86 | 0.75 |

For each of our degraded RNA preparations, mean increases in Ct for housekeeping and non-housekeeping genes did not differ (t-tests; 0.25 < p < 0.40). Furthermore, regression analysis revealed strong correlations (r > 0.999; Fig. 3) between the increases in housekeeping and non-housekeeping C_t_ among the six degraded RNA preparations, spanning a wide range of mean C_t_ increases corresponding to levels of RNA degradation from approximately 50% to more than 95%.

**Fig. 3.**
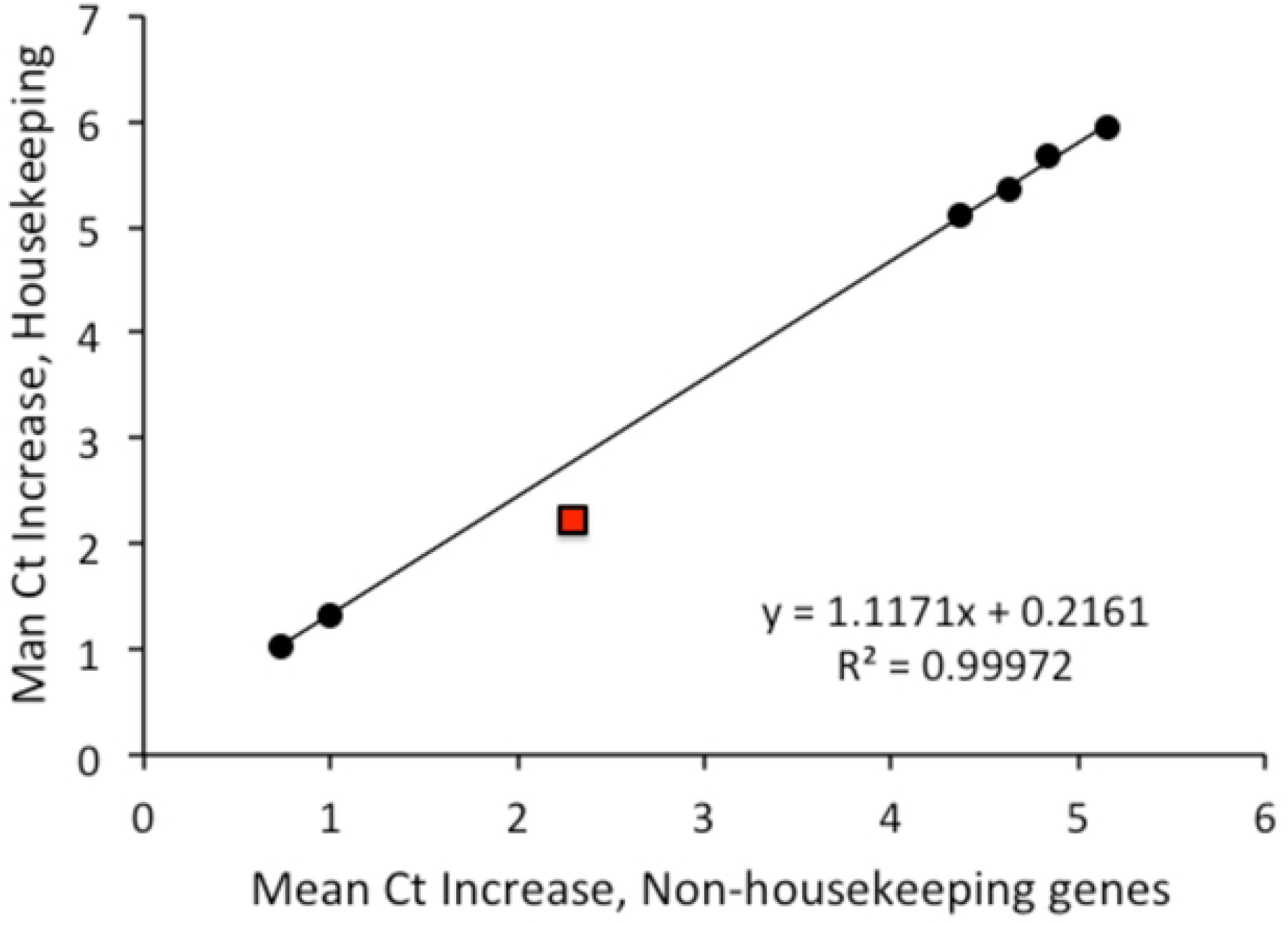
Regression analysis of relative RNA degradation effects on mean C_t_ values for housekeeping and non-housekeeping genes among the 89 targets evaluated. Note the strong linear relationship spanning a range of C_t_ increases from approximately one cycle in the least degraded preparations (corresponding to ∼50% mean loss of target mRNAs) to approximately six cycles in the most highly degraded preparations (corresponding to >95% mean loss of target mRNAs). Black data points are results from *in vitro* RNA digestions, and the red data point is from *in situ* RNA degradation in frozen and thawed ACL (see next).

To determine if RNase digestion of RNA *in vitro* accurately modeled the *in situ* degradation of RNA in poorly stored tissues, we thawed ACL tissue to permit action of endogenous RNases prior to RNA isolation, and compared the gene expression profile from the degraded RNA to that obtained with intact RNA. Thawing 20 min before initiating extraction produced overt degradation, with decreased amounts of intact rRNAs and increased amounts of smaller digestion products (Fig. 4A) in comparison to RNA from unthawed ACL. Nevertheless, as observed for *in vitro* digested RNAs, the gene expression profile obtained using RNA from thawed ACL correlated strongly with the profile from unthawed ACL (R^2^ = 0.92). Furthermore, *in situ* digestion produced a slope of the degraded-vs.-intact regression line (0.915, Fig. 4B) very close to slope of 0.906 from the *in vitro* digestion experiments. Finally, *in situ* digestion produced very similar losses of housekeeping and non-housekeeping gene mRNAs (mean Ct increases of 2.2 and 2.3 respectively, reflecting more than 75% loss of amplifiable targets in the tissue thawed for 20 min), comparable (red data point in Fig. 3) to the relative losses observed in the *in vitro* digestion experiments.

**Fig. 4.**
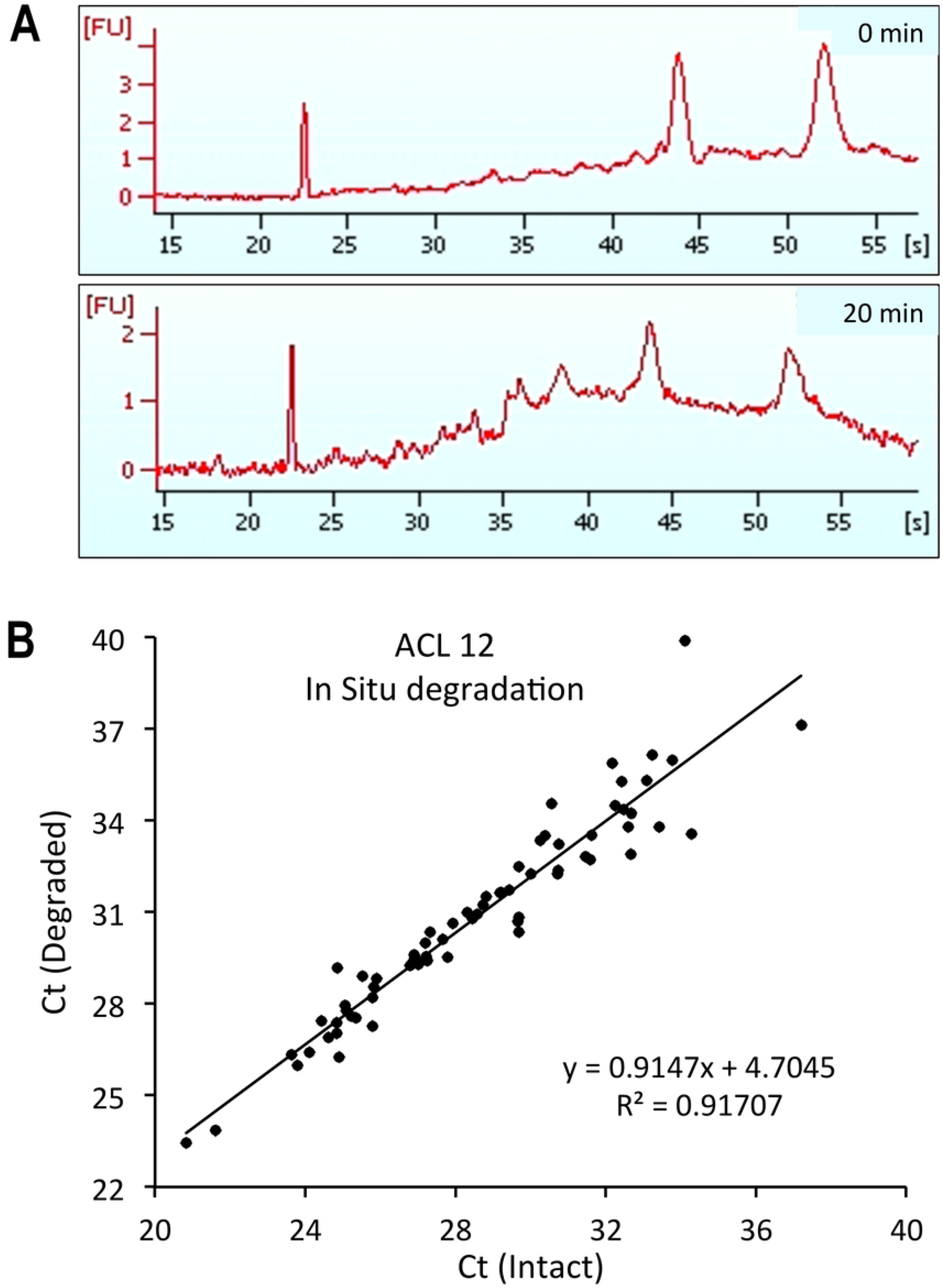
Comparison of expression profiles from intact and *in situ*-degraded RNA. Panel A shows Bioanalyzer traces of RNA isolated 0 min (upper trace) or 20 min (lower trace) after thawing the ACL tissue at 23°C. The peaks at 44 and 52 min are ribosomal RNAs. For the RNA isolated 20 min post-thaw, note the increased baseline and smaller rRNA peaks, reflecting the autolytic degradation of RNAs to smaller products. Panel B shows the corresponding linear regression plot of Ct values from the intact vs. *in situ*-degraded RNAs. Note the strong correlation (R^2^ = 0.92) between the profiles and the very similar slope of the regression line (0.91) to that observed in the *in vitro* digestion experiments (Table 1).

## DISCUSSION

Here we found that partial degradation of human ACL RNA decreased the amounts of individual mRNAs amplified in RT-PCR-based expression analyses but did not dramatically alter the overall gene expression profile in comparison to intact RNA. We also found no significant difference between the loss of housekeeping and loss of non-housekeeping mRNAs within degraded RNA preparations. Therefore, by normalizing to levels of housekeeping mRNAs, legitimate comparisons can be made between profiles of mRNAs with varying levels of degradation. Indeed, up to 97% mean loss of amplifiable target still yielded profiles that correlated strongly with profiles obtained from intact RNA, albeit with loss of signals from some low abundance targets. Further digestion of RNA would be required to define the point at which excessive degradation of ACL RNA yields invalid expression profiles. Regardless, our results revealed that valid, PCR-based expression profiles can be generated from RNA that is much more highly degraded than is generally thought to be required for such studies.

ACL RNA digested *in vitro* with RNase ONE™ could differ from RNA degraded by endogenous cellular ribonucleases. RNase ONE™ is a broad specificity enzyme from *E. coli* that cleaves between any two ribonucleotides, whereas human cellular ribonucleases exhibit more restricted nucleobase specificities^12^. Nevertheless, differences in enzyme action are unlikely to affect our findings for two reasons. First, the promiscuous activity of RNase ONE™ theoretically approximates the combined specificities of the multiple RNases^15^ in human cells that together mediate *in situ* RNA degradation in a poorly handled or stored tissue. Second, hydrolysis of an mRNA anywhere between the upstream PCR primer sequence and the downstream priming site for reverse transcriptase would render it incapable of serving as a template for RT-PCR. Consequently, any endogenous RNase, including one that preferentially hydrolyzed at certain ribonucleotides (such as the CpX specificity of pancreatic RNase A ^16^), would be expected to digest target mRNAs sufficiently to cause loss of RT-PCR signal much as we achieved by digestion with RNase ONE™. Indeed, expression analysis of RNA degraded *in situ* in frozen-and-thawed ACL confirmed the conceptual soundness of the *in vitro* digestion approach in this study.

Our results do not rule out the possibility that certain individual RNAs might be highly sensitive to hydrolysis by specific RNases; however, use of RNA that is presumed to be completely intact also does not necessarily preclude that possibility. Of note, by using a combination of oligo-dT and random hexamers to prime the RT step for cDNA synthesis, we maximized yields of first strand template for downstream amplification by PCR. Thus, even RNA preparations degraded entirely to relatively small fragments could still generate expression signals so long as individual mRNA fragments both spanned their corresponding PCR-amplified target sequences and extended far enough downstream to provide a site for oligo-dT or random hexamer priming. Accordingly, the results of this study do show that useful expression information for most genes can be obtained from human connective tissues acquired in a way that does not assure complete integrity of RNA (e.g., from biopsies or cadaveric specimens).

Although RNA in poorly stored or handled tissues will ultimately degrade to the point that valid expression data cannot be obtained, we conclude it is nevertheless possible to generate meaningful results from partially degraded RNA so long as sufficient mRNA template remains for PCR amplification. This finding is particularly relevant to research correlating gene expression profiles with biomechanical properties of cadaveric tissues, including studies on sex differences in molecular and biomechanical properties of the human ACL, and on age-and pathology-associated differences in the molecular and biomechanical properties of intervertebral discs.

## ACKNOWLEDGEMENTS

We thank Elizabeth White for outstanding technical assistance. DMH and JRS received grant number AR049767 from the National Institute for Arthritis and Musculoskeletal and Skin Diseases (https://www.niams.nih.gov/), and DMH and JH received funding from the TTU/TTUHSC Clinical/Basic Science Seed Grant Program. The funders had no role in study design, data collection and analysis, decision to publish, or preparation of the manuscript.

